# Transcription drives DNA replication initiation and termination in human cells

**DOI:** 10.1101/324079

**Authors:** Yu-Hung Chen, Sarah Keegan, Malik Kahli, Peter Tonzi, David Fenyö, Tony T. Huang, Duncan J. Smith

## Abstract

The locations of active DNA replication origins in the human genome, and the determinants of origin activation, remain controversial. Additionally, neither the predominant sites of replication termination nor the impact of transcription on replication-fork mobility have been defined. We demonstrate that replication initiation occurs preferentially in the immediate vicinity of the transcription start site of genes occupied by high levels of RNA polymerase II, ensuring co-directional replication of the most highly transcribed genes. Further, we demonstrate that dormant replication origin firing represents the global activation of pre-existing origins. We also show that DNA replication naturally terminates at the polyadenylation site of transcribed genes. During replication stress, termination is redistributed to gene bodies, generating a global reorientation of replication relative to transcription. Our analysis provides a unified model for the coupling of transcription with replication initiation and termination in human cells.

## INTRODUCTION

Since the identification of *cis*-acting sequences responsible for the definition of replication origins in *S. cerevisiae* (Stinchcomb et al., 1979), significant effort has been applied to identify analogous determinants of replication initiation in human cells (Hyrien, 2015; Prioleau and MacAlpine, 2016). However, although origins have been observed via several independent techniques to be enriched close to transcribed genes and a range of other chromatin features (Besnard et al., 2012; Dellino et al., 2013; Langley et al., 2016; Petryk et al., 2016), a coherent model that encompasses both origin specification and activation has not emerged. Throughout eukaryotes, many more replication origins are licensed by MCM2-7 loading in G1 than are required to complete S-phase (Donovan et al., 1997; Edwards et al., 2002). The pool of excess MCMs is required for survival when dNTPs are depleted by the ribonucleotide reductase inhibitor hydroxyurea (HU) (Ge et al., 2007), and has been proposed to allow the firing of ‘dormant’ replication origins that rescue genome replication after replication fork stalling. The identities of these dormant origins, and how they differ from constitutive origins, have not been defined.

DNA is a one-dimensional template that can be simultaneously acted upon by the replication and transcription machineries. The orientation of essential genes in prokaryotes is biased to avoid head-on collisions between these two processes (Rocha, 2003); among eukaryotes, budding yeast and *C. elegans* show statistical orientation bias of the most highly transcribed genes to the co-directional orientation (Osmundson et al., 2017; Pourkarimi et al., 2016), and significant co-orientation of transcribed genes has been noted in human cells (Hamperl et al., 2017; Petryk et al., 2016). Head-on replication-transcription conflicts are deleterious in eukaryotes, leading to increased DNA damage (Hamperl et al., 2017) and genomic rearrangements (Tran et al., 2017). The mechanisms by which origin location is specified to co-orient replication with transcription in diverse human cell types, and the impact of co-directional transcription on replication-fork progression through genes, remain speculative.

Here, by using Okazaki fragment sequencing (Ok-seq) to infer the direction of replication-fork movement, we define transcription initiation efficiency and gene length as independent determinants of replication origin location and firing efficiency, and show that origin firing occurs close to the transcription start site (TSS), ensuring co-oriented replication of the most highly transcribed genes. We additionally show that dormant origins correspond to constitutive origins. Further, we determine the existence of widespread replication-fork stalling due to co-oriented replication-transcription conflicts at transcription termination sites (TTS) under unperturbed conditions, and in gene bodies during replication stress.

## RESULTS

Using Ok-seq, we and others have reported on replication initiation (McGuffee et al., 2013; Petryk et al., 2016; Pourkarimi et al., 2016), replication elongation (Osmundson et al., 2017) and lagging-strand processing (Osmundson et al., 2017; Smith and Whitehouse, 2012). To investigate replication initiation and termination in untransformed human cells, we performed Ok-seq (Petryk et al., 2016) on hTERT-immortalized RPE-1 cells. Previous genome-wide studies of metazoan replication show limited agreement in origin calls (Hyrien, 2015; Prioleau and MacAlpine, 2016), with the exception that all identify significant enrichment of origins close to transcribed genes. Therefore, instead of aiming to identify all potential sites at which replication can possibly initiate in the human genome, we focused our analysis on the efficiency of replication initiation and termination in transcribed regions.

### Replication initiates in the immediate vicinity of high-conflict transcription start sites

Rightward-moving replication forks generate Okazaki fragments (OFs) that map to the Crick strand, while leftward-moving forks generate Watson-strand fragments (Fig. 1A). Therefore, a replication origin will manifest as an increase in the proportion of OFs mapping to the Crick strand: the efficiency and spatial localization of origin firing will impact the magnitude and gradient of this transition, respectively (Fig. 1B). OFs showed no strand bias around random genomic loci (Fig. S1). Consistent with global origin activity at TSS, meta-analysis of OF Crick strand bias over a 50 kb window surrounding all 17920 annotated TSS showed a symmetrical transition from predominantly leftward-to predominantly rightward-moving forks (Fig. 1C). However, separate analysis of Watson-and Crick-strand genes revealed a profound asymmetry based on gene orientation (Fig. 1D). Therefore, for all subsequent analyses we considered OF strand bias relative to gene orientation, such that transcription occurs from left to right and OF polarity is computationally reversed for Crick-strand genes. All data for TSS analysis were highly reproducible across two biological replicates (Fig. S2).

**Figure 1.**
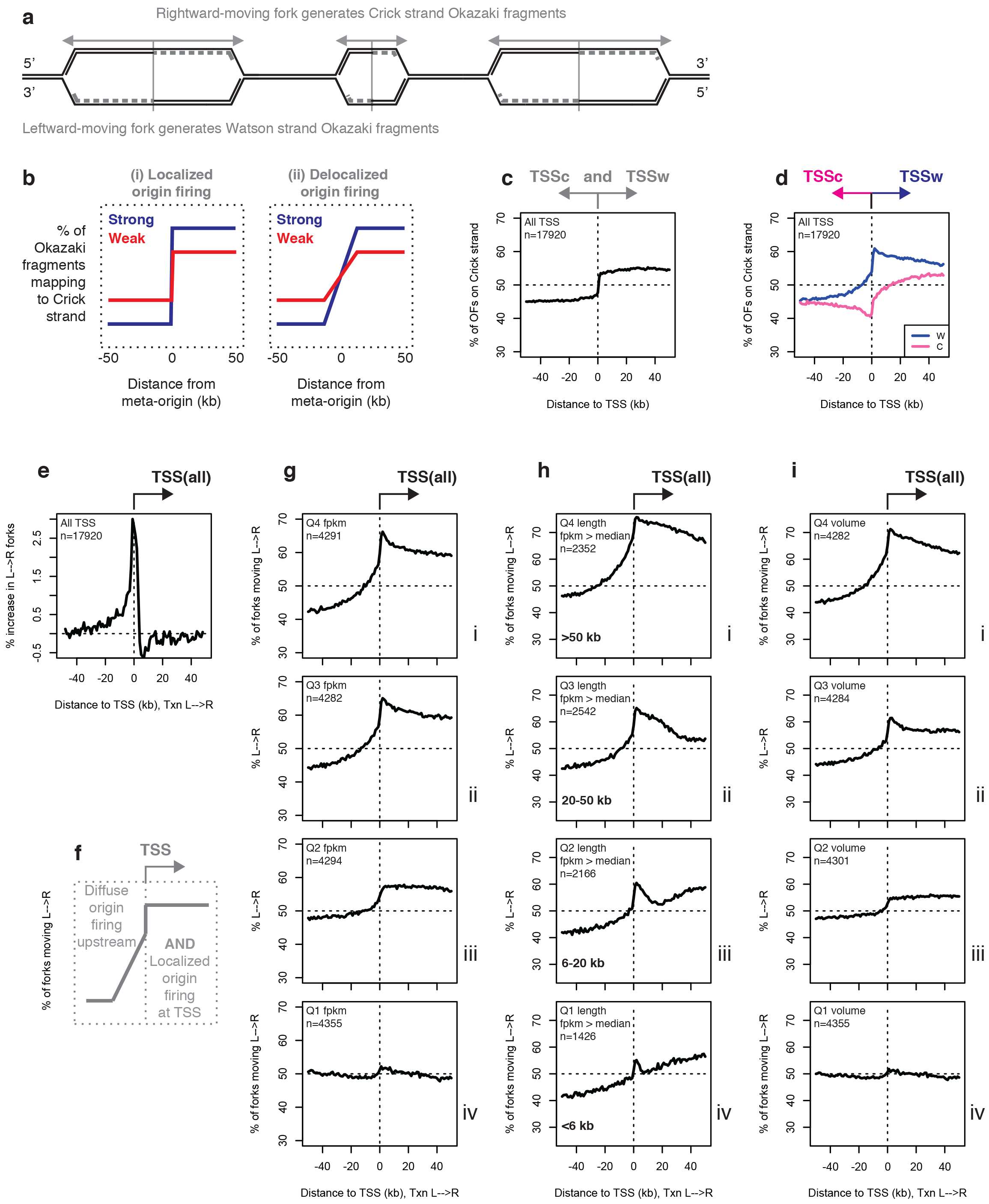
Replication initiates preferentially at the TSS of long, transcribed genes: transcriptional volume predicts replication origin efficiency. **(A)** The relationship between replication direction and Okazaki fragment (OF) strand. **(B)** Expected OF distributions around replication origins. **(C)** Percent of OFs mapping to the Crick strand across a ±50 kb window around all transcription start sites (TSS) in the human genome. All data in figures 1-3 are from asynchronously dividing hTERT-immortalized RPE-1 cells, and are displayed using 1 kb bins. **(D)** Percentage of OFs mapping to the Crick strand around the TSS of Watson (W) or Crick (C) genes. **(E)** First derivative of Okazaki fragment strand bias around TSS, oriented such that transcription occurs from left to right. Data are smoothed across two 1 kb bins. **(F)** Schematic representation of upstream and TSS-proximal replication initiation inferred from data in Figure 1. **(G)** Percentage of replication forks moving left to right around TSS binned by RNA-seq read depth quartile from (Harenza et al., 2017) (panels i-iv). **(H)** Percentage of replication forks moving left to right around TSS of actively transcribed genes (FPKM > median, (Harenza et al., 2017)) binned by length according to quartiles for all genes (panels i-iv). **(I)** Percentage of replication forks moving left to right around TSS binned by transcriptional volume (FPKM from (Harenza et al., 2017) x gene length).

By analyzing the first derivative of replication direction, we can directly infer the extent of replication initiation at each position relative to the meta-TSS. We observe that, while individually low but cumulatively high levels of initiation occur over a wide region upstream of TSS, initiation is strongly biased to a ~5 kb region immediately adjacent to the TSS itself and is under-represented or absent in gene bodies (Fig. 1E, schematic in Fig. 1F).

To investigate the effect of transcription on TSS-proximal replication initiation, we separated genes into quartiles based on RNA-seq read density (FPKM, Fragments Per Kilobase of transcript per Million mapped reads) across the gene body in RPE-1 cells (Harenza et al., 2017). TSS of genes with higher FPKM showed significantly greater change in strand bias than TSS of weakly or non-transcribed genes (Fig. 1G). However, we also observed a significant length dependence in this effect, such that short genes have weaker origin activity than long genes with equivalent RNA-seq read density (Fig. 1H). Gene length and FPKM are not correlated in RPE-1 cells (r = −0.044): therefore, gene length and transcript number independently correlate with origin firing efficiency. While (assuming equal RNA decay rates) FPKM reports the number of RNA molecules synthesized, the number of RNA polymerases occupying a gene for a given FPKM is linearly related to the length of the gene i.e. a 50 kb gene producing one transcript per minute will be occupied by ten times as many polymerases as a similarly transcribed 5 kb gene. We therefore analyzed OF strand bias around TSS separated by transcriptional volume (FPKM x gene length). Origin activity was strongly correlated with high transcriptional volume (Fig. 1I). All data from Fig. 1 were reproducible in previously published Ok-seq data from HeLa cells (Petryk et al., 2016) (Fig. S3). We note that the distance to the nearest downstream TSS or transcription termination site (TTS) - both of which are intrinsically dependent on gene length - are substantially stronger predictors of origin efficiency than the distance to the nearest upstream TSS or TTS, which are independent of gene length (Fig. S4). Therefore, the effect of gene length on origin efficiency does not simply reflect decreased passive replication from nearby ‘competitor’ origins. We conclude that, in unperturbed human cells, the initiation of DNA replication is strongly biased towards the immediate vicinity of TSS that drive transcription of genes with high RNA polymerase II (RNAP2) occupancy. Thus, while transcription inherently creates conflicts with replication, the coupling of origin firing to RNAP2 occupancy ensures that these conflicts are in the co-directional orientation.

### Modulation of constitutive origin efficiency during replication stress

In response to replication stress, origin firing increases: this is generally interpreted as the activation of dormant origins (Blow et al., 2011). However, firing events during growth in hydroxyurea (HU) have previously been reported to occur in the vicinity of transcribed genes (Karnani and Dutta, 2011). To analyze origin firing efficiency at TSS in the context of increased or decreased dormant origin usage, we treated cells with 0.2 mM HU for 4h before OF collection, and additionally depleted either of the Fanconi Anemia (FA) effector proteins, (Ceccaldi et al., 2016; Michl et al., 2016) FANCD2 or FANCI, by RNAi. This HU treatment regime has previously been shown to decrease average inter-origin distance without activating the checkpoint (Ge et al., 2007): we previously showed that knockdown of FANCI increases inter-origin distance in HU, reflecting reduced origin firing, while knockdown of FANCD2 weakly stimulates origin firing under the same conditions (Chen et al., 2015). In the absence of HU, knockdown of either FANCI or FANCD2 have little or no measureable effect on origin firing, cell cycle progression, or cell doubling (Chen et al., 2015). OF distributions around TSS showed a greater change in strand bias adjacent to high-volume TSS for HU-treated cells relative to untreated (Fig. 2A), consistent with further increased firing efficiency of efficient origins. The rapid decrease in strand bias downstream of the TSS can be explained by increased intragenic fork stalling and is addressed below (Fig. 3). Knockdown of FANCI in HU-treated cells reduced strand bias around high-volume TSS, while knockdown of FANCD2 very slightly increased it (Fig. 2B & S2C). Both short and long genes were affected (Fig. S5). Therefore, under conditions of globally increased or decreased origin firing, we find that the firing efficiency of the most active origins is the most strongly affected. Our data suggest that intragenic replication origins are not required for widespread dormant origin firing, and that dormant origins are in fact constitutive origins with altered firing efficiency.

**Figure 2.**
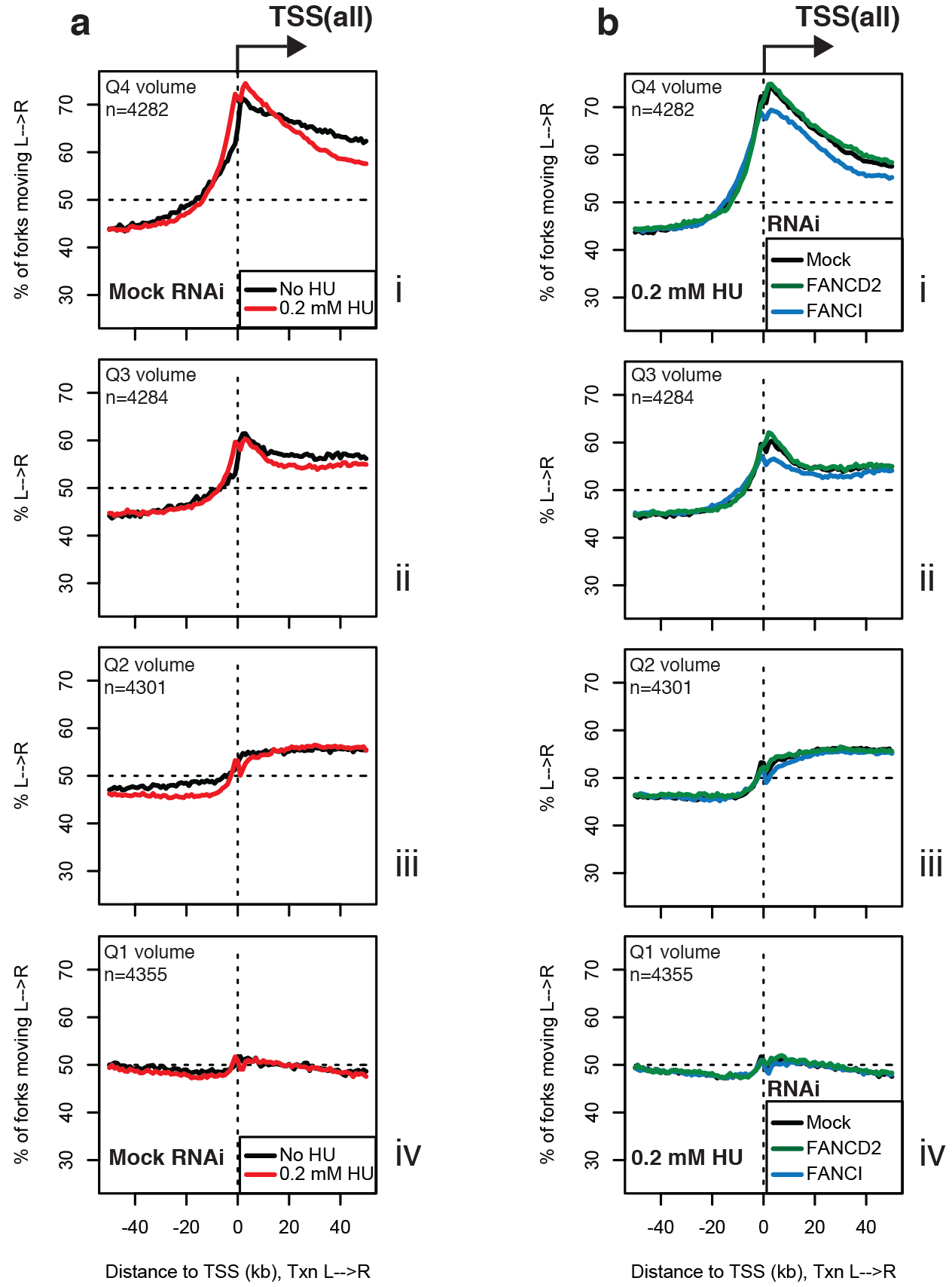
Global modulation of origin activity under conditions that increase or suppress dormant origin firing. **(A)** Percentage of replication forks moving left to right around TSS binned by transcriptional volume, for cells grown in the absence (black) or presence (red) of 0.2 mM hydroxyurea (HU) for 4h before OF collection. **(B)** Percentage of replication forks moving left to right around TSS binned by transcriptional volume, for cells treated with siRNAs against FANCD2 (green), FANCI (blue), or mock-treated (black), grown in 0.2 mM hydroxyurea for 4h before OF collection.

### Replication terminates at transcription termination sites

We reasoned that the loss of OF strand bias observed at a gene-length-dependent distance downstream of the TSS (Fig. 1H) might result from replication termination at or near the 3’ ends of transcribed genes. To test this, we queried OF distributions around the annotated gene 3’ end (hereafter referred to as the TTS). The TTS corresponds to the site of pre-mRNA cleavage and polyadenylation, and may not directly reflect the site of RNAP2 dissociation for genes that rely on Xrn2-mediated transcriptional termination (Proudfoot, 2016). The number of RNA polymerases terminating transcription should equal the number initiating at the TSS; therefore, we analyzed OF strand bias around the TTS of genes separated by quartiles based on FPKM, which serves as a proxy for initiation frequency. Consistent with widespread replication termination at the TTS of transcribed genes, we observed a dramatic, transcription-dependent reduction in replication forks moving in the direction of transcription through the gene body, occurring precisely at the TTS (Fig. 3A). This termination signal was surrounded by an increase in left-to-right polarity, consistent with diffuse origin firing occurring a variable distance away at nearby TSS (Fig. 3B).

**Figure 3.**
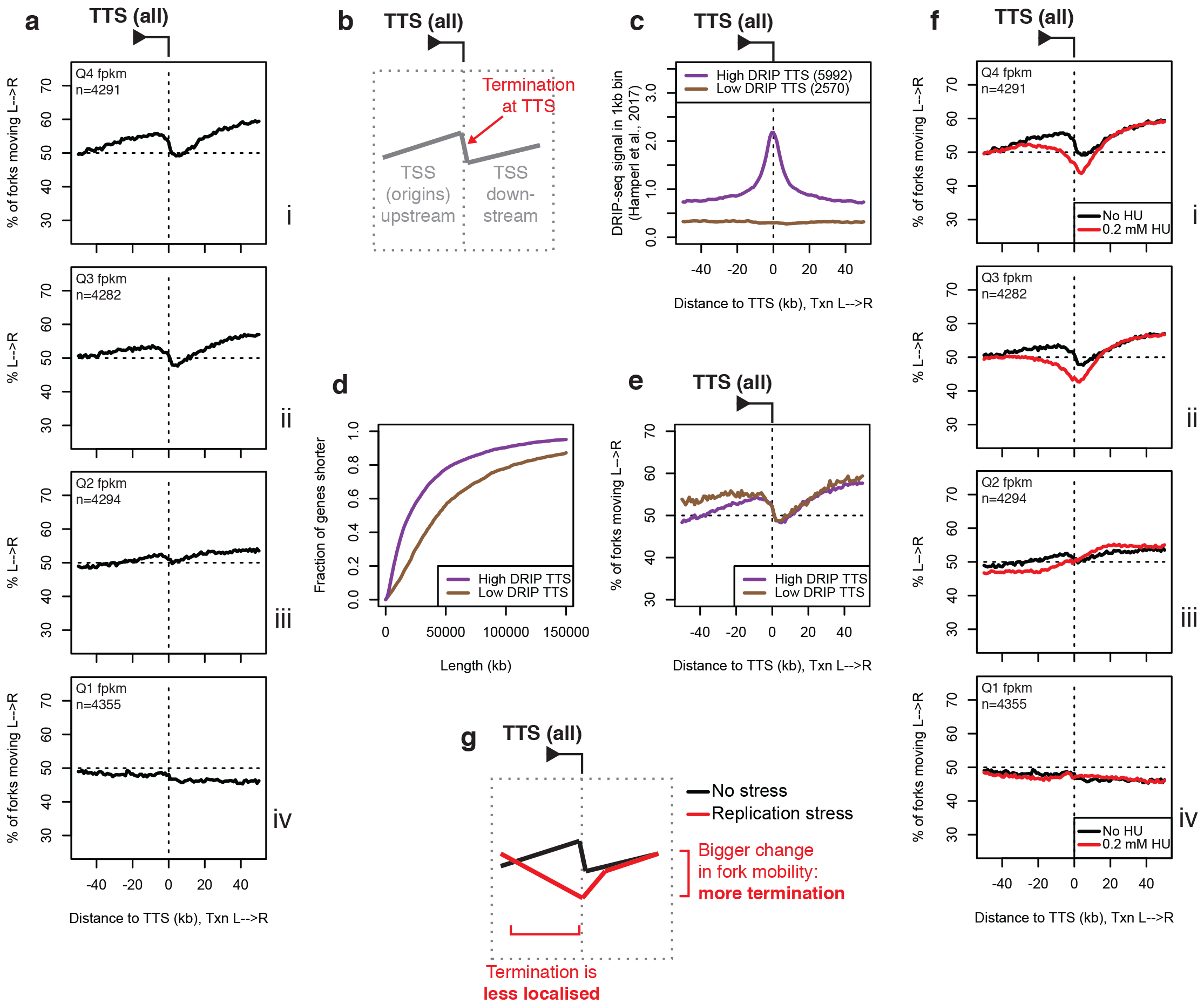
Widespread, R-loop-independent replication-fork termination occurs at the 3’ ends of transcribed genes under unperturbed conditions. Termination in gene bodies increases under replication stress. **(A)** Percentage of replication forks moving left to right around transcription termination sites (TTS) binned by RNA-seq read depth quartile from (Harenza et al., 2017) (panels i-iv). **(B)** Schematic representation of replication termination at TTS, with initiation at proximal TSS up-and downstream. **(C)** DRIP-seq signal from (14) around TTS of actively transcribed genes (FPKM from (18) > median) separated into high (purple) and low (brown) DRIP bins based on DRIP-seq signal (Hamperl et al., 2017) ±10kb from the TTS. **(D)** Length distribution of actively transcribed high-DRIP vs low-DRIP genes **(E)**Percentage of replication forks moving left to right around TTS of actively transcribed (FPKM from (Harenza et al., 2017) > median) high-DRIP vs low-DRIP genes. **(F)(A)** Percentage of replication forks moving left to right around (TTS) binned by RNA-seq read depth quartile from (Harenza et al., 2017) (panels i-iv), for cells grown in the absence (black) or presence (red) of 0.2 mM hydroxyurea (HU) for 4h before OF collection. **(G)**Schematic representation of the change in replication termination observed under replication stress.

Because RNA-DNA hybrids (R-loops) have been proposed to modulate both replication-fork stalling and DNA damage, we analyzed genes separately based on their likely levels of R-loop formation at TTS in RPE cells. Most differences in R-loop formation between cell types and even species can be attributed to differences in transcription (Sanz et al., 2016): therefore, we separated genes with FPKM above the median level in RPE cells into two bins based on the DRIP-seq signal observed ±10 kb from their TTS in HeLa cells (Hamperl et al., 2017), to obtain high-and low-DRIP TTS gene sets (Fig. 3C). We note that the genes in our high-DRIP set are significantly shorter than those in our low-DRIP set (Fig. 3D), possibly reflecting some contribution of signal from TSS-proximal R-loops in short genes. Analysis of OF strand bias around the TTS of high-and low-DRIP genes indicated equivalent replication termination regardless of the propensity to form R-loops (Fig. 3E). The upward gradient observed upstream of the TTS for high-DRIP genes is due to origin firing at TSS, which are closer to TTS due to the length difference observed in Fig. 3D. Data from Fig. 3 A-E were reproducible in previously published Ok-seq data from HeLa cells (Petryk et al., 2016) (Fig. S6), and are therefore not attributable to differences in R-loop formation between cell types. Thus, we conclude that replication naturally terminates at the 3’ end of transcribed genes, and that termination is independent of R-loop formation. Replication origins are located at variable distances from TTS: therefore, this effect must be driven by replication-fork stalling or arrest as opposed to simply by replication initiation kinetics (McGuffee et al., 2013).

We next analyzed replication-fork mobility around TTS in the context of growth in 0.2 mM HU (Fig. 3F, schematic in Fig. 3G). All data for TTS were again reproducible across replicate datasets (Fig. S7). We observed both an increase in the overall level of gene-associated replication termination, and a change in the predominant location of termination from the TTS to the gene body. Consistent with this observation, analysis of OF strand bias around TSS indicates that the proportion of replication forks moving co-directionally through gene bodies decreases more rapidly in the presence than the absence of 0.2 mM HU (Fig. 2A), but is unaffected by knockdown of FANCI or FANCD2 (Fig. 2B). Thus, replication termination is significantly altered by 0.2 mM HU: we propose that replication forks normally stall upon encountering RNAP2 paused at the TTS, but that intragenic replication-transcription conflicts become the predominant cause of replication-fork stalling under replication stress.

## DISCUSSION

### Co-orientation of replication and transcription in multicellular eukaryotes

Our data suggest that DNA replication initiation is inherently coupled to transcription initiation, such that replication preferentially initiates immediately adjacent to the TSS of genes with high RNAP2 occupancy during S-phase (model, Fig. S8). Spatially limiting initiation in transcribed genomic regions to the TSS of the most highly transcribed genes ensures that these show the strongest bias towards co-directional replication in any proliferative cell type, regardless of its transcriptional profile. Activation of additional genes, for example during differentiation or oncogenesis (Macheret and Halazonetis, 2018), would concurrently change the replication profile of the cell to mitigate conflicts with transcription. Many unicellular organisms, including prokaryotes and *S. cerevisiae* define replication origins via the use of *cis-acting* sequences: in these organisms, the orientation of conflict-prone genes has been biased by evolution (Merrikh, 2017; Osmundson et al., 2017; Paul et al., 2013), presumably as a result of damage induced by head-on collisions followed by re-orientation (Srivatsan et al., 2010). However, the use of such *cis*-acting sequences in an organism with many transcriptionally distinct proliferative cell types would be problematic because it would enforce deleterious head-on replication-transcription conflicts in a cell-type-specific fashion. Thus, while unicellular organisms can co-orient transcription with replication by genome evolution, the alternative strategy - to co-orient replication with transcription - is more robust for organisms with many distinct transcriptomes.

A simple mechanism to explain the increased replication origin activity of the genes occupied by the most RNAP2 is that chromatin accessibility independently determines both MCM2-7 loading and the recruitment of replisome components to these licensed replication origins. We note that chromatin accessibility, as opposed to RNA-seq read density, has previously been reported as the best correlate of replication timing (Hansen et al., 2009). Under conditions of global origin activation or repression, a proportionally similar activation/repression of all origins will lead to the greatest change in origin efficiency occurring at origins with intrinsically high firing efficiencies (Fig. 2). Thus, dormant origin firing globally maintains co-orientation of replication and transcription, and can simply be considered as a change in average origin efficiency as opposed to the specific regulation of distinct classes of origin.

There are fewer actively transcribed genes than the number of origins required to replicate the human genome. Inactive regions of the genome lack both spatially localized regions of highly accessible chromatin and a requirement to co-orient replication with transcription: therefore, specifically localized replication origins would serve no biological purpose in these regions. Transient chromatin opening, likely driven by low-levels of pervasive transcription factor binding and/or transcription, could ensure a sufficient density of MCM2-7 loading and activation to support genome duplication in these regions through a distribution of origin firing events that largely disfavors initiation at specific individual sites. We speculate that the poor overlap between genome-wide origin mapping studies (Prioleau and MacAlpine, 2016) is due to a lack of localized origin firing outside highly transcribed regions as opposed to an intrinsic flaw with any origin detection approach.

Intuitively, replication-fork initiation at TSS followed by stalling or arrest at TTS might impair the replication of intragenic regions flanked by convergent, highly transcribed genes. However, the ability of both RNA polymerases (Gros et al., 2014; Gros et al., 2015) and the replicative helicase (Douglas et al., 2018) to push loaded MCM2-7 double hexamers would lead to a gradual redistribution of un-fired origins towards these intragenic regions, where they could serve as a pool of licensed ‘rescue’ origins.

### Re-orientation of replication-transcription conflicts during replication stress

Although DNA replication and transcription are, on average, co-oriented throughout the actively transcribed regions of the genome, we observe a robust replication termination signal at the TTS of transcribed genes (Fig. 3). Therefore, co-oriented collisions between the replication and transcription machineries are highly prevalent, and RNAP paused at the 3’ ends of genes acts as a physical barrier to even co-directional replication-fork progression. Replication-fork stalling due to co-oriented collisions with RNA polymerase at gene 3’ ends has previously been observed in prokaryotes (Mirkin et al., 2006) but not, to our knowledge, in eukaryotes. Unlike in prokaryotes (Lang et al., 2017), but similarly to replication-fork arrest at tRNA genes in yeast (Osmundson et al., 2017), R-loops do not appear to impede replisome mobility despite their impact on the ultimate outcome of collisions (Tran et al., 2017; Hamperl et al., 2017; Paulsen et al., 2009; Stirling et al., 2011). We note that independent screens in HeLa cells (Paulsen et al., 2009) and *S. cerevisiae* (Stirling et al., 2011) identified RNA splicing and 3’ end formation factors as preventative against DNA damage. It will be interesting to determine the relative contributions of R-loop modulation and replisome mobility to the increased damage when RNA processing is impaired.

The locations of replication origins are minimally affected by hydroxyurea-mediated dNTP depletion, but the progression of replication forks through genes is severely impeded (Fig. 2A, 3F). We propose that, during replication stress, co-directional replication-transcription conflicts within the gene body become more frequent due to reduced replisome speed, leading to intragenic replisome stalling. Under these conditions, replication of gene 3’ ends must be rescued by a head-on fork. Such a global re-orientation of replication-transcription conflicts towards the head-on orientation would greatly increase DNA damage and ATR signaling, which are strongly orientation-dependent (Hamperl et al., 2017). Therefore, we propose that conflict re-orientation, as opposed to unscheduled dormant origin firing or impaired replication fork movement *per se* due to replication-transcription conflicts, underlies the toxic effects of replication stress and the transcription-dependent fragility of long genes (Helmrich et al., 2011).

## ACKNOWLEDGEMENTS

We thank the NYU Genome Technology Center for assistance with TapeStation and sequencing. We thank D. Remus, I. Whitehouse, E. Mazzoni, and H. Klein for helpful discussions. Y-H C was funded in part by the Molecular Oncology and Immunology NCI training program through NYU School of Medicine (5T32CA009161-40). Work in T.H. laboratory is supported by grants from the NIH (ES025166), V Foundation BRCA Research and Basser Innovation Award. Work in D.J.S. laboratory is supported by grants from the NIH (GM127336, GM114340) and the Searle Scholars Program.

## DATA AVAILABILITY

A token to access the raw and processed data is available upon request from the corresponding authors.

## AUTHOR CONTRIBUTIONS

Y-H. C., M.K. and P.T. performed research, S.K. and D.J.S. analyzed data., D.F., T.T.H. and D.J.S. conceived and supervised the study. All authors interpreted data. D.J.S. wrote the manuscript with input from all authors.

## METHODS

### Cell Culture/EdU labeling

hTERT immortalized RPE-1 cells (ATCC) were grown in Dulbecco’s modified Eagle’s medium: Nutrient Mixture F-12 (DMEM/F-12) media (Life Technologies) supplemented with 10% fetal bovine serum (FBS), 3% sodium bicarbonate and 1% Pen-Strep. In brief, exponentially growing cells of 50-60% confluency were plated in 150mm dishes (approximately 15-20 plates per sample/replicate). siRNA transfections were done according to (Chen et al., 2015). EdU labeling was done for 2 min (4 min for low-dose HU) at 20 μM final concentration. Cells were either untreated or treated with low-dose HU (200 μM final concentration).

### Ok-seq

Okazaki fragments were purified and libraries generated essentially as described (Petryk et al., 2016), with the most major modification being the use of gel-purified adaptor duplexes from (Smith and Whitehouse, 2012). Libraries were sequenced using the HiSeq-2500 platform and reads were aligned to the hg19 build of the human genome.

### Data analysis

TSS and TTS locations were obtained from the UCSC genome browser http://genome.ucsc.edu - genes devoid of uniquely mapping OF reads on both the Watson and Crick strand through the entire gene body were excluded from analysis. Genes were separated by RNA-seq read density in RPE-1 cells, length or DRIP-seq read density in HeLa cells as described in the main text.

---

**Figure S1.**
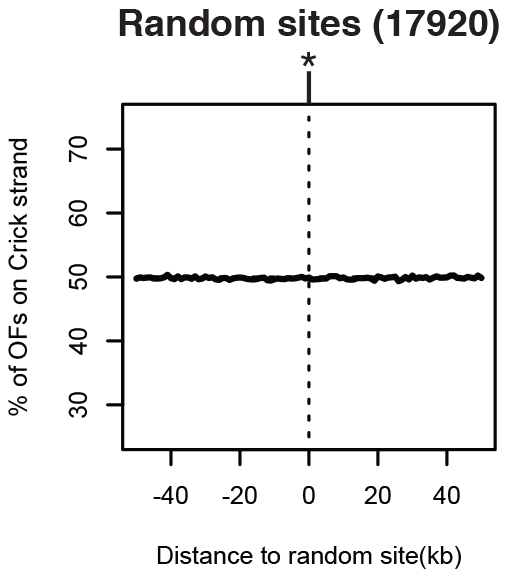
Okazaki fragments show no strand bias around random genomic loci.

**Figure S2.**
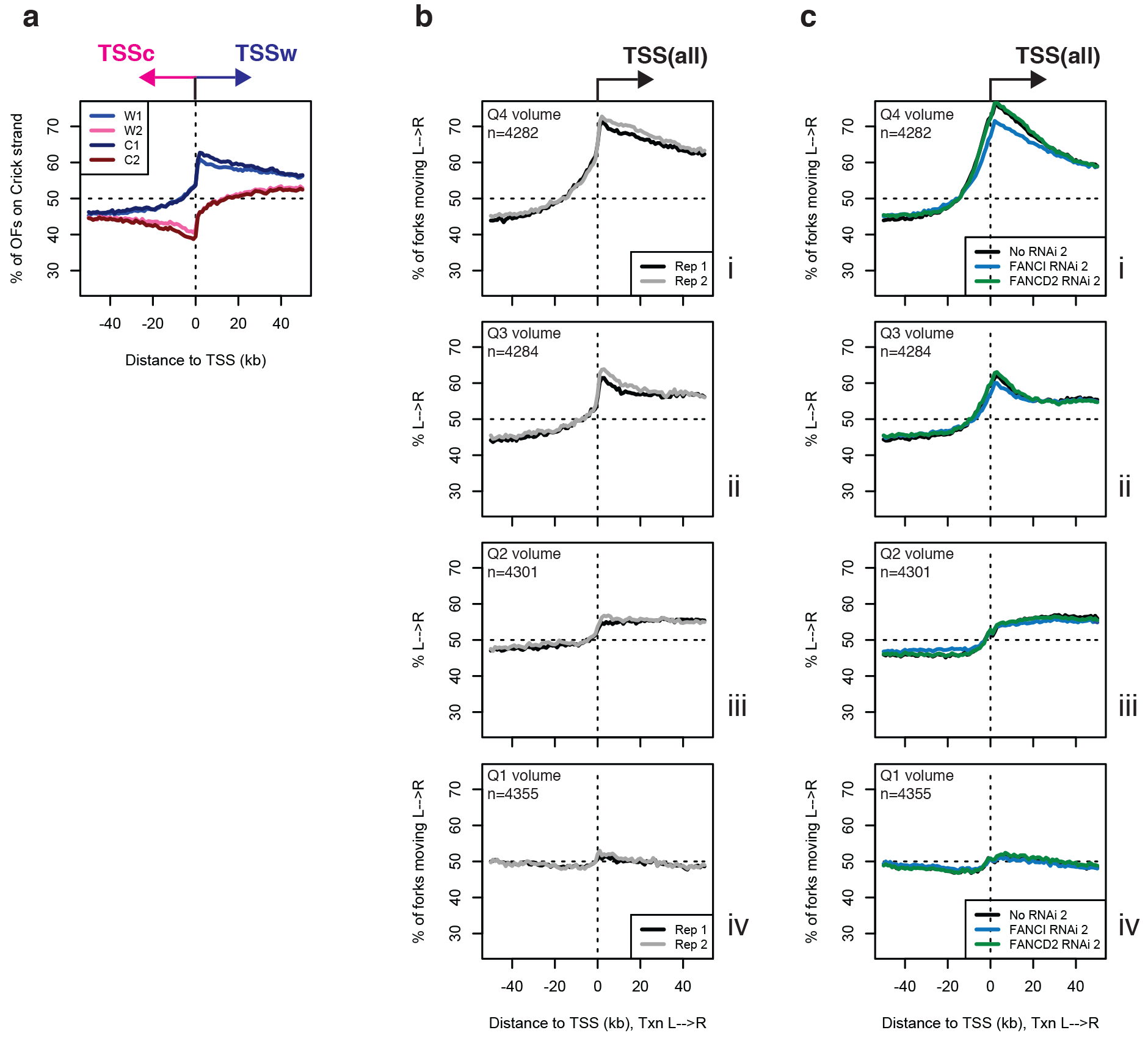
Reproducibility of TSS data across replicate datasets. **(A)** Data were analyzed as in Fig. 1B, for two replicate datasets. **(B)** Data were analyzed as in Fig. 1G, for two replicate datasets. **(C)** Data were analyzed as in Fig.2B, using the second replicate datasets for each knockdown condition.

**Figure S3.**
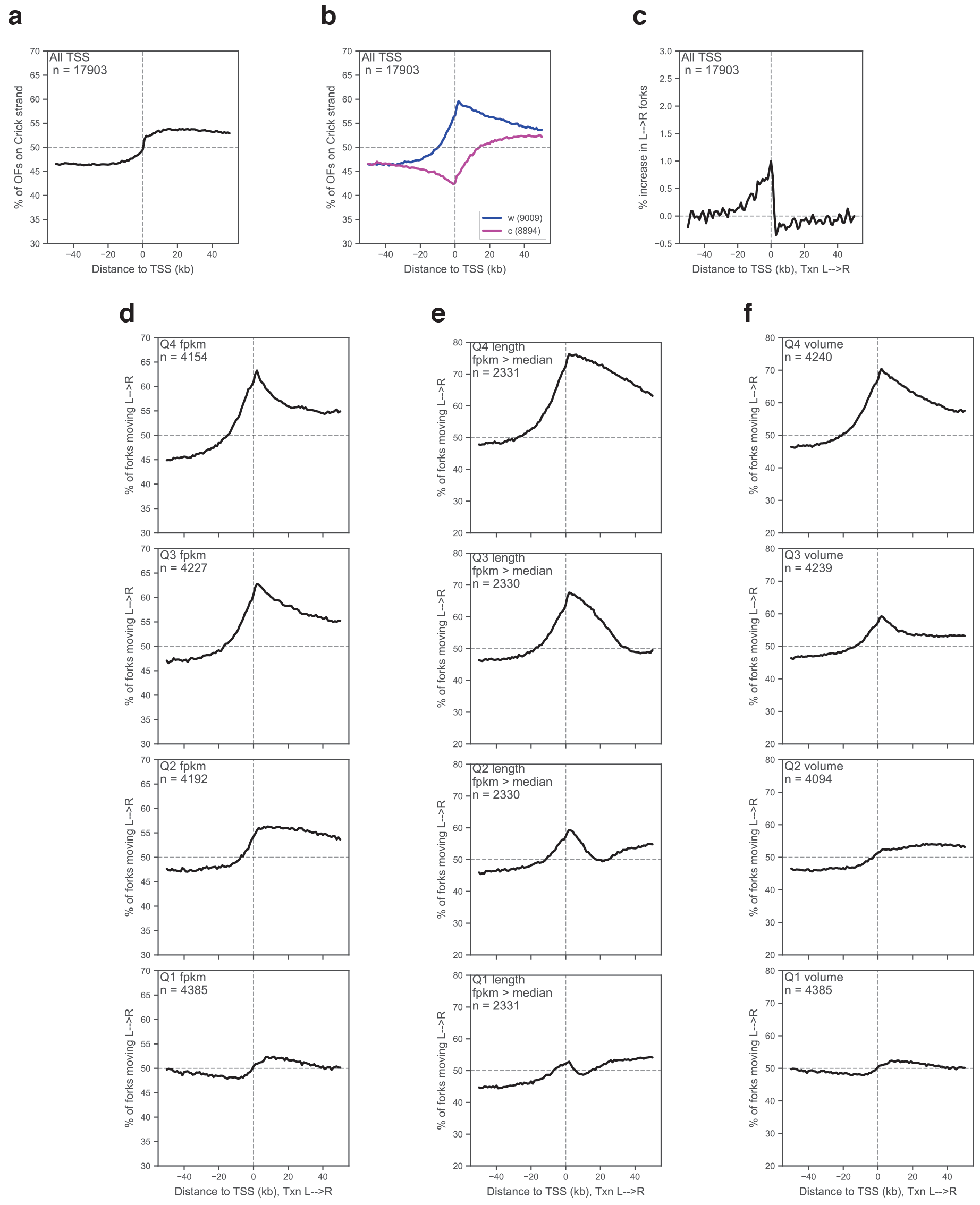
Replication initiation is most efficient at high-volume TSS in HeLa cells. **(A)** Data were analyzed as in Fig. 1C, using Ok-seq data from HeLa cells (7) **(B)** Data were analyzed as in Fig. 1D, using Ok-seq data from HeLa cells (7) **(C)** Data were analyzed as in Fig. 1E, using Ok-seq data from HeLa cells (7) **(D)** Data were analyzed as in Fig. 1G, using Ok-seq data from HeLa cells (7) **(E)** Data were analyzed as in Fig. 1H, using Ok-seq data from HeLa cells (7) **(F)** Data were analyzed as in Fig. 1I, using Ok-seq data from HeLa cells (7) All gene expression data for HeLa cells are from the EBI gene expression atlas https://www.ebi.ac.uk/gxa/home - absent values for FPKM were set to zero.

**Figure S4.**
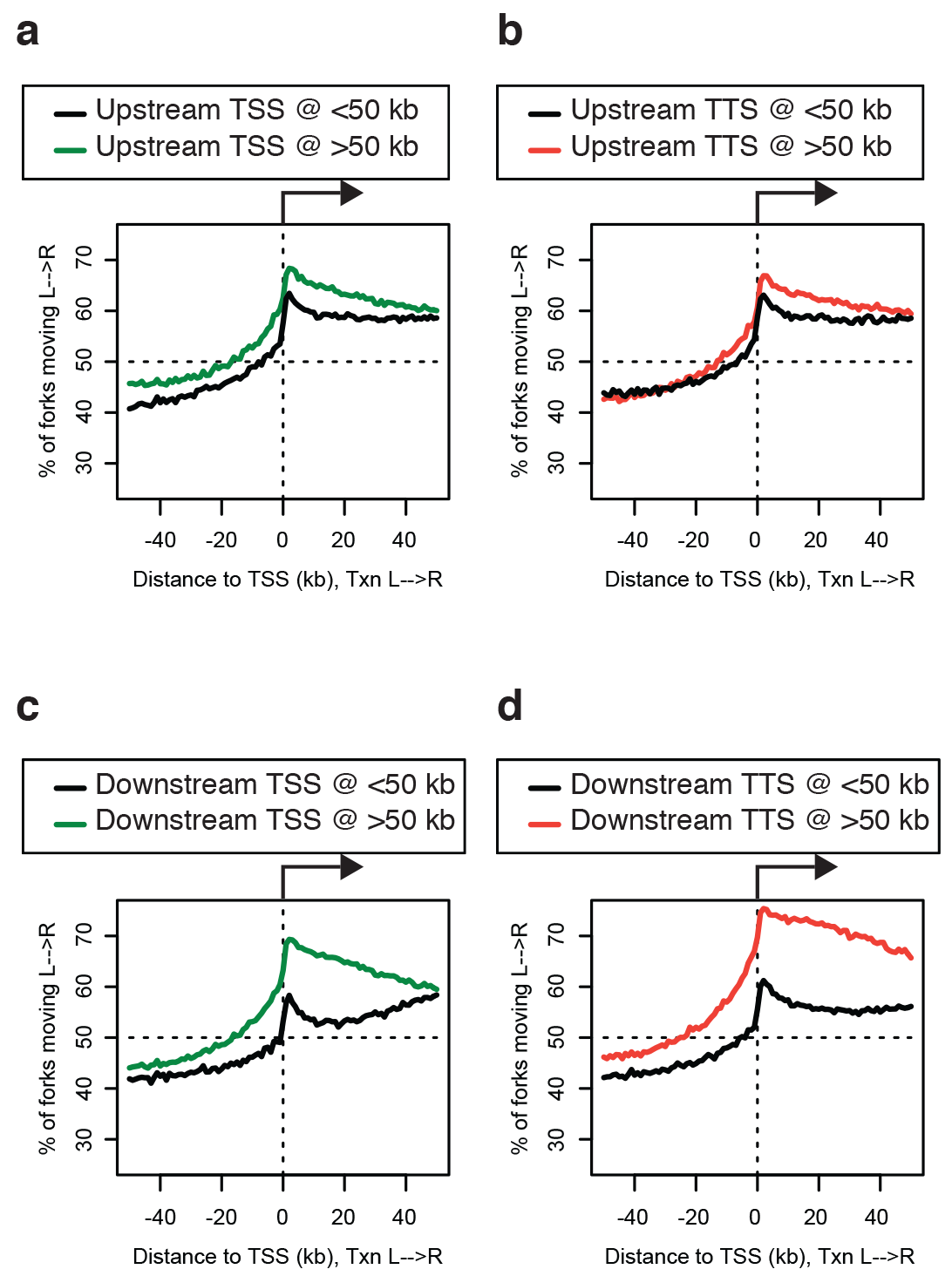
The effect of gene length on TSS-proximal origin firing efficiency is not solely a result of passive replication. **(A)** Percentage of replication forks moving left to right around TSS of actively transcribed genes (FPKM (18) > median), where the TSS of the most proximal upstream gene is under (black) or over (green) 50 kb from the TSS being analyzed. **(B)** Percentage of replication forks moving left to right around TSS of actively transcribed genes (FPKM (18) > median), where the TTS of the most proximal upstream gene is under (black) or over (red) 50 kb from the TSS being analyzed. **(C)** Percentage of replication forks moving left to right around TSS of actively transcribed genes (FPKM (18) > median), where the TSS of the most proximal downstream gene is under (black) or over (green) 50 kb from the TSS being analyzed. **(D)** Percentage of replication forks moving left to right around TSS of actively transcribed genes (FPKM (18) > median), where most proximal downstream TTS is under (black) or over (red) 50 kb from the TSS being analyzed. Note that this TSS-TTS distance is equivalent to the length of the gene.

**Figure S5.**
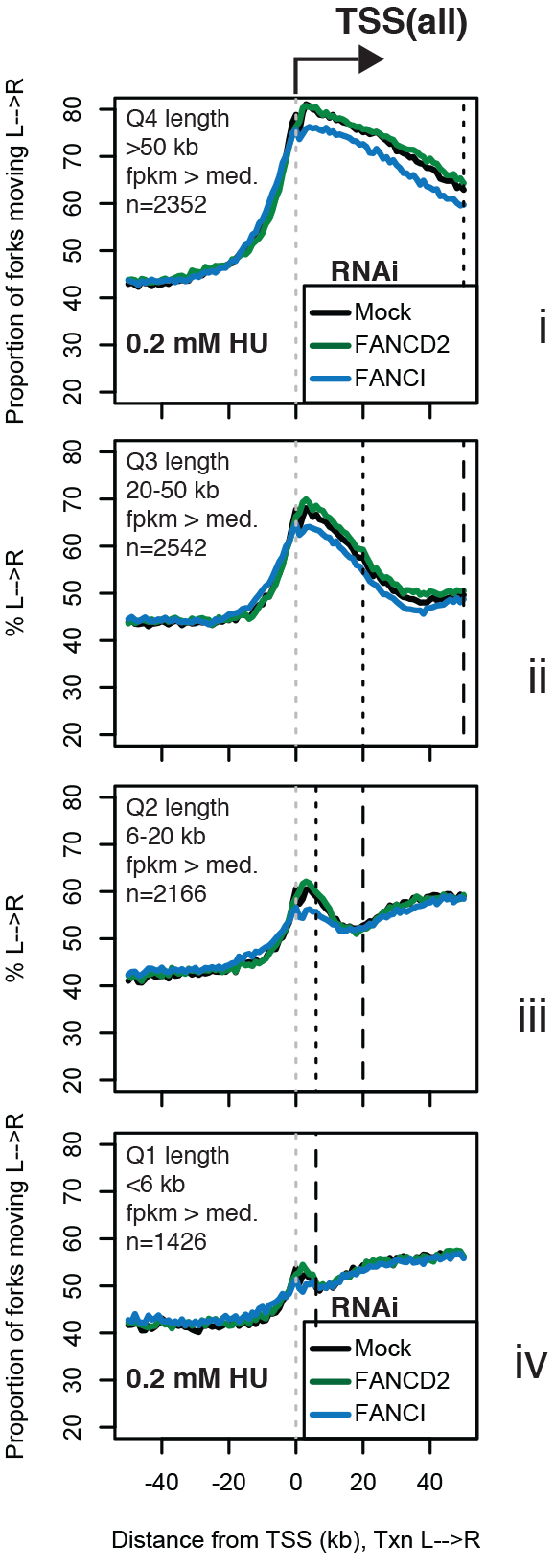
The effect of FANCI or FANCD2 knockdown is not related to gene length or increased replication termination. Percentage of replication forks moving left to right around TSS binned by transcriptional volume (FPKM (18) × gene length) for cells treated with siRNAs against FANCD2 (green), FANCI (blue), or mock-treated (black), grown in 0.2 mM hydroxyurea for 4h before OF collection.

**Figure S6.**
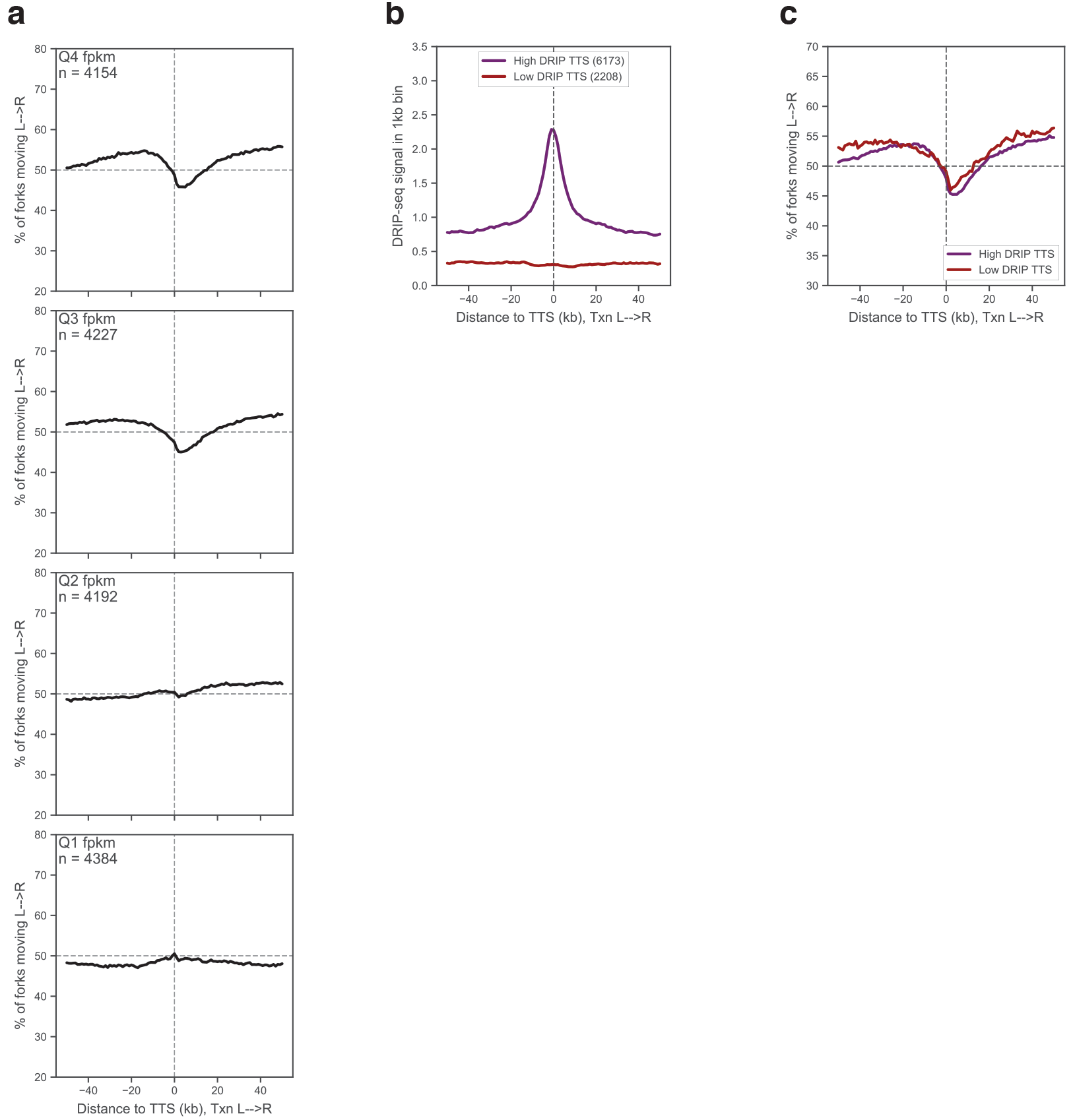
Transcription-dependent, R-loop-independent replication termination at TTS in HeLa cells. **(A)** Data were analyzed as in Fig. 3A, using Ok-seq data from HeLa cells (7) **(B)** Data were analyzed as in Fig. 3C: DRIP-seq data are from HeLa cells (14) **(C)** Data were analyzed as inFig. 3E, using Ok-seq data from HeLa cells (7): : DRIP-seq data are from HeLa cells (14)

**Figure S7.**
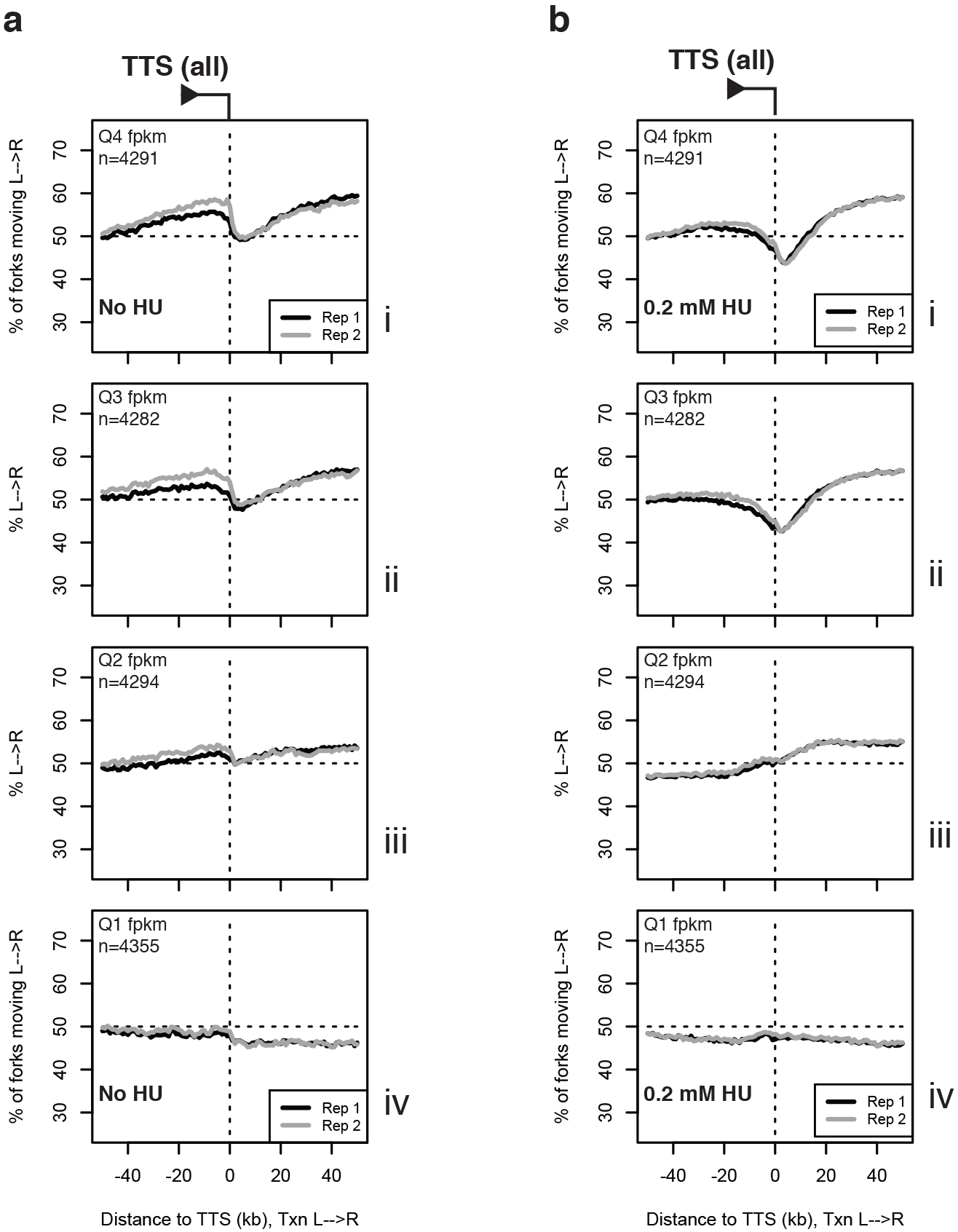
Reproducibility of TTS data across replicate datasets. **(A)** Data were analyzed as in Fig. 3A, for two replicate datasets. **(B)** Data from cells grown in 0.2 mM HU for 4h before OF collection were analyzed as in Fig. 3F, for two replicate datasets.

**Figure S8.**
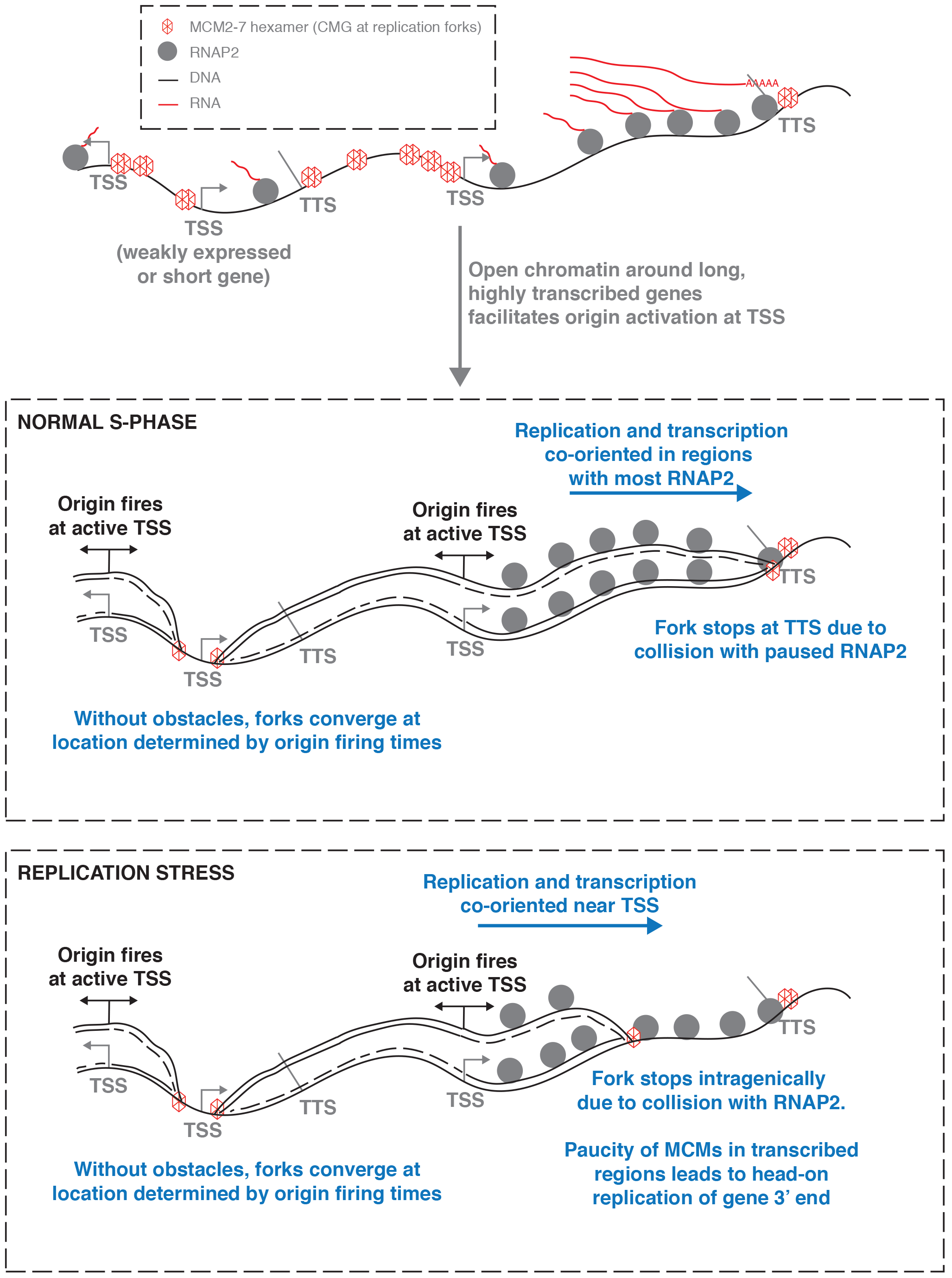
Model.

## REFERENCES

Besnard, E., Babled, A., Lapasset, L., Milhavet, O., Parrinello, H., Dantec, C., Marin, J. M., and Lemaitre, J. M. (2012). Unraveling cell type-specific and reprogrammable human replication origin signatures associated with G-quadruplex consensus motifs. Nat Struct Mol Biol 19, 837–844.

Blow, J. J., Ge, X. Q., and Jackson, D. A. (2011) How dormant origins promote complete genome replication. Trends Biochem Sci 36, 405–414.

Ceccaldi, R., Sarangi, P., and D’Andrea, A. D. (2016) The Fanconi anaemia pathway: new players and new functions. Nat Rev Mol Cell Biol 17, 337–349.

Chen, Y. H., Jones, M. J., Yin, Y., Crist, S. B., Colnaghi, L., Sims, R. J., Rothenberg, E., Jallepalli, P. V., and Huang, T. T. (2015) ATR-mediated phosphorylation of FANCI regulates dormant origin firing in response to replication stress. Mol Cell 58, 323–338.

Dellino, G. I., Cittaro, D., Piccioni, R., Luzi, L., Banfi, S., Segalla, S., Cesaroni, M., Mendoza-Maldonado, R., Giacca, M., and Pelicci, P. G. (2013) Genome-wide mapping of human DNA-replication origins: levels of transcription at ORC1 sites regulate origin selection and replication timing. Genome Res 23, 1–11.

Donovan, S., Harwood, J., Drury, L. S., and Diffley, J. F. (1997). Cdc6p-dependent loading of Mcm proteins onto pre-replicative chromatin in budding yeast. Proc Natl Acad Sci U S A 94, 5611–5616.

Douglas, M. E., Ali, F. A., Costa, A., and Diffley, J. F. X. (2018) The mechanism of eukaryotic CMG helicase activation. Nature 555, 265–268.

Edwards, M. C., Tutter, A. V., Cvetic, C., Gilbert, C. H., Prokhorova, T. A., and Walter, J. C. (2002) MCM2-7 complexes bind chromatin in a distributed pattern surrounding the origin recognition complex in Xenopus egg extracts. J Biol Chem 277, 33049–33057.

Ge, X. Q., Jackson, D. A., and Blow, J. J. (2007) Dormant origins licensed by excess Mcm2-7 are required for human cells to survive replicative stress. Genes Dev 21, 3331–3341.

Gros, J., Devbhandari, S., and Remus, D. (2014) Origin plasticity during budding yeast DNA replication in vitro. EMBO J 33, 621–636.

Gros, J., Kumar, C., Lynch, G., Yadav, T., Whitehouse, I., and Remus, D. (2015) Post-licensing Specification of Eukaryotic Replication Origins by Facilitated Mcm2-7 Sliding along DNA. Mol Cell 60, 797–807.

Hamperl, S., Bocek, M. J., Saldivar, J. C., Swigut, T., and Cimprich, K. A. (2017) Transcription-Replication Conflict Orientation Modulates R-Loop Levels and Activates Distinct DNA Damage Responses. Cell 170, 774–786.e19.

Hansen, R. S., Thomas, S., Sandstrom, R., Canfield, T. K., Thurman, R. E., Weaver, M., Dorschner, M. O., Gartler, S. M., and Stamatoyannopoulos, J. A. (2009). Sequencing newly replicated DNA reveals widespread plasticity in human replication timing. Proc Natl Acad Sci U S A 107, 139–144.

Harenza, J. L., Diamond, M. A., Adams, R. N., Song, M. M., Davidson, H. L., Hart, L. S., Dent, M. H., Fortina, P., Reynolds, C. P., and Maris, J. M. (2017) Transcriptomic profiling of 39 commonly-used neuroblastoma cell lines. Sci Data 4, 170033.

Helmrich, A., Ballarino, M., and Tora, L. (2011) Collisions between replication and transcription complexes cause common fragile site instability at the longest human genes. Mol Cell 44, 966–977.

Hyrien, O. (2015) Peaks cloaked in the mist: the landscape of mammalian replication origins. J Cell Biol 208, 147–160.

Karnani, N., and Dutta, A. (2011) The effect of the intra-S-phase checkpoint on origins of replication in human cells. Genes Dev 25, 621–633.

Lang, K. S., Hall, A. N., Merrikh, C. N., Ragheb, M., Tabakh, H., Pollock, A. J., Woodward, J. J., Dreifus, J. E., and Merrikh, H. (2017) Replication-Transcription Conflicts Generate R-Loops that Orchestrate Bacterial Stress Survival and Pathogenesis. Cell 170, 787–799.e18.

Langley, A. R., Gräf, S., Smith, J. C., and Krude, T. (2016) Genome-wide identification and characterisation of human DNA replication origins by initiation site sequencing (ini-seq). Nucleic Acids Res 44, 10230–10247.

Macheret, M., and Halazonetis, T. D. (2018) Intragenic origins due to short G1 phases underlie oncogene-induced DNA replication stress. Nature 555, 112–116.

McGuffee, S. R., Smith, D. J., and Whitehouse, I. (2013) Quantitative, Genome-Wide Analysis of Eukaryotic Replication Initiation and Termination. Mol Cell 50, 123–135.

Merrikh, H. (2017) Spatial and Temporal Control of Evolution through Replication-Transcription Conflicts. Trends Microbiol 25, 515–521.

Michl, J., Zimmer, J., and Tarsounas, M. (2016) Interplay between Fanconi anemia and homologous recombination pathways in genome integrity. EMBO J 35, 909–923.

Mirkin, E. V., Castro Roa, D., Nudler, E., and Mirkin, S. M. (2006). Transcription regulatory elements are punctuation marks for DNA replication. Proc Natl Acad Sci U S A 103, 7276–7281.

Osmundson, J. S., Kumar, J., Yeung, R., and Smith, D. J. (2017) Pif1-family helicases cooperatively suppress widespread replication-fork arrest at tRNA genes. Nat Struct Mol Biol 24, 162–170.

Paul, S., Million-Weaver, S., Chattopadhyay, S., Sokurenko, E., and Merrikh, H. (2013) Accelerated gene evolution through replication-transcription conflicts. Nature 495, 512–515.

Paulsen, R. D., Soni, D. V., Wollman, R., Hahn, A. T., Yee, M. C., Guan, A., Hesley, J. A., Miller, S. C., Cromwell, E. F., Solow-Cordero, D. E., Meyer, T., and Cimprich, K. A. (2009) A genome-wide siRNA screen reveals diverse cellular processes and pathways that mediate genome stability. Mol Cell 35, 228–239.

Petryk, N., Kahli, M., d’Aubenton-Carafa, Y., Jaszczyszyn, Y., Shen, Y., Silvain, M., Thermes, C., Chen, C. L., and Hyrien, O. (2016) Replication landscape of the human genome. Nat Commun 7, 10208.

Pourkarimi, E., Bellush, J. M., and Whitehouse, I. (2016) Spatiotemporal coupling and decoupling of gene transcription with DNA replication origins during embryogenesis in C. elegans. Elife 5,

Prioleau, M. N., and MacAlpine, D. M. (2016) DNA replication origins-where do we begin. Genes Dev 30, 1683–1697.

Proudfoot, N. J. (2016) Transcriptional termination in mammals: Stopping the RNA polymerase II juggernaut. Science 352, aad9926.

Rocha, E. P. C. (2003) Gene essentiality determines chromosome organisation in bacteria. Nucleic Acids Research 31, 6570–6577.

Sanz, L. A., Hartono, S. R., Lim, Y. W., Steyaert, S., Rajpurkar, A., Ginno, P. A., Xu, X., and Chédin, F. (2016) Prevalent, Dynamic, and Conserved R-Loop Structures Associate with Specific Epigenomic Signatures in Mammals. Mol Cell 63, 167–178.

Smith, D. J., and Whitehouse, I. (2012) Intrinsic coupling of lagging-strand synthesis to chromatin assembly. Nature 483, 434–438.

Srivatsan, A., Tehranchi, A., MacAlpine, D. M., and Wang, J. D. (2010) Co-orientation of replication and transcription preserves genome integrity. PLoS Genet 6, e1000810.

Stinchcomb, D. T., Struhl, K., and Davis, R. W. (1979) Isolation and characterisation of a yeast chromosomal replicator. Nature 282, 39–43.

Stirling, P. C., Bloom, M. S., Solanki-Patil, T., Smith, S., Sipahimalani, P., Li, Z., Kofoed, M., Ben-Aroya, S., Myung, K., and Hieter, P. (2011) The complete spectrum of yeast chromosome instability genes identifies candidate CIN cancer genes and functional roles for ASTRA complex components. PLoS Genet 7, e1002057.

Tran, P. L. T., Pohl, T. J., Chen, C. F., Chan, A., Pott, S., and Zakian, V. A. (2017) PIF1 family DNA helicases suppress R-loop mediated genome instability at tRNA genes. Nat Commun 8, 15025.

